# Regulation of NFKBIZ by precise Regnase-1 endoribonuclease cleavage and subsequent uridylation

**DOI:** 10.1101/2025.09.19.677199

**Authors:** Samira Ebrahimi, Richard Zapf, Isabelle Gracien, Krysta Koyle, Ryan Morin, Timothy Eric Audas, Peter J. Unrau

**Affiliations:** Department of Molecular Biology and Biochemistry, Simon Fraser University, Burnaby, British Columbia, Canada V5A 1S6

**Author notes:** To whom communication should be addressed.

**Keywords:** Regnase-1, NFKBIZ, endoribonuclease, uridylation, Post Transcriptional Regulation, RNA Degradation, COVID-19, DLBCL, immune response

## Abstract

A conserved sequence in the 3′UTR of NFKBIZ mRNA has long been recognized as a regulator of cytokine production and interferon responses. We show that the endoribonuclease Regnase-1 controls NFKBIZ expression through a precise and modular RNA degradation mechanism. The structured core element undergoes specific endonucleolytic cleavage, while flanking upstream and downstream stem–loop modules, previously implicated in Regnase-1 recognition, act cooperatively to enhance cleavage efficiency by ∼25-fold. Following cleavage, the upstream fragment is rapidly uridylated, accelerating decay of the NFKBIZ open reading frame. This pathway explains how driver mutations – found in this RNA region – responsible for diffuse large B-cell lymphoma elevate NFKBIZ expression and how a segment of the SARS-CoV-2 genome – previously linked to NFKBIZ activation – suppresses Regnase-1 cleavage via hybridization to this regulatory RNA segment. Together, these findings define a mechanistic framework for Regnase-1–mediated control of NFKBIZ, linking its cleavage activity to both lymphomagenesis and viral pathogenesis.

## INTRODUCTION

Inflammation is a fundamental host defense mechanism that protects against microbial pathogens and environmental stressors, while also contributing to cancer progression and viral infection^1,2^. Although essential for activating adaptive immunity, uncontrolled or unresolved inflammation can drive autoimmune disease, cytokine storms, and chronic inflammatory disorders^3–6^. Maintaining immune homeostasis requires precise regulation of inflammatory responses, achieved through multiple levels of gene control. Among these, post-transcriptional regulation plays a central role in fine-tuning cytokine expression^7^. Such pathways, often mediated by conserved ribonucleases^8^, are frequently disrupted in inflammatory disease ^9^, subverted during viral infection ^10,11^, or circumvented by cancer-associated mutations^12,13^.

Regnase-1 (Zc3h12a, MCPIP1) is a conserved endoribonuclease within a family of four proteins (Regnase-1 to 4), all of which contain a CCCH-type zinc-finger domain and a catalytic RNase center ^14^. Mice lacking Regnase-1 usually die within twelve weeks, as they develop severe anemia, uncontrolled plasma cell expansion, and systemic inflammation ^15^. Predominantly expressed in immune cells, Regnase-1 regulates Toll-like receptor signaling in response to lipopolysaccharide and related stimuli^16,17^ by destabilizing transcripts encoding pro-inflammatory cytokines (e.g., TNF, IL-1β, IL-12b, and IL-6, IL-17), immune regulators (NFKBIZ, NFKBID) and its own mRNA^18–21^. Expression peaks within two hours of immune stimulation, consistent with a tightly regulated feedback loop^22^. After cleavage, targeted transcripts are known to undergo rapid decay, but the molecular mechanisms underlying endonuclease and degradation processes remain poorly defined.

IκBζ, which is encoded by NFKBIZ mRNA, is one of the most established targets of Regnase-1 (REF). This NF-κB transcriptional co-factor, selectively modulates a subset of NF-κB-dependent genes, thereby playing a pivotal role in inflammatory signaling ^23^. Regulation of NFKBIZ by Regnase-1 has been attributed to UPF1 helicase dependent cleavage of a conserved 3′UTR sequence immediately downstream of the stop codon^24,25^. This region coincides with recurrent driver mutations in diffuse large B-cell lymphoma (DLBCL)^26^ and can hybridize with a SARS-CoV-2 RNA segment, stabilizing NFKBIZ during infection^27^.

Here, we delineate a molecular mechanism by which Regnase-1 specifically recognizes and cleaves the NFKBIZ 3′UTR, initiating rapid mRNA degradation. We identify a core cleavage motif comprising three highly conserved stem-loops, with additional upstream and downstream elements that enhance cleavage efficiency by ∼25-fold. Following cleavage, the upstream fragment undergoes extensive TUT7-dependent uridylation, incorporating up to 34 U residues as revealed by high-throughput sequencing. This polyuridylated fragment is then degraded by the exonuclease DIS3L2, as confirmed by RNAi knockdown. Together, our findings establish a coupled Regnase-1–TUT7–DIS3L2 pathway as the central mechanism governing NFKBIZ regulation, directly linking Regnase-1 activity to both lymphomagenesis and viral pathogenesis.

## RESULTS

### Identification of a precise Regnase-1-dependent RNA cleavage motif within the 3∼ UTR of NFKBIZ

The 3′ untranslated region (UTR) of NFKBIZ is 1,651 nucleotide long, and contains a highly conserved region (from human to Zebrafish) of 184 nucleotide surrounding the stop codon that we call the TNU (**T**runcated **N**FKBIZ 3′ **U**TR) (**Figure 1A and S1**). Chemical probing and T1 RNase mapping experiments suggest that the secondary structure of the TNU consists of seven conserved stem-loop structures^19,26^ labeled here as SL0 to SL6 (**Figure 1B**) that appeared likely targets for Regnase-1 cleavage. To identify the precise Regnase-1 cleavage site we incubated 5′ ^32^P end-labeled TNU with purified Regnase-1, and found a time-dependent RNA cleavage site in the loop region SL2, between nucleotides C_59_ and A_60_ (**Figure 1B and C**). Minor secondary and tertiary cleavage sites were also detected between SL3 and SL4, occurring immediately after nucleotides G_100_ and C_102_, but these were more than 100-fold weaker than the predominant cleavage observed after C_59_ (**Figure 1C**). Further characterization of dominant cleavage products revealed that Regnase-1 produces an upstream 5′ fragment possessing a terminal 3′ hydroxyl and a downstream fragment possessing a 5′ phosphate, consistent with a water-mediated, magnesium-dependent cleavage mechanism (**Figure S2 and S3**).

**Figure 1.**
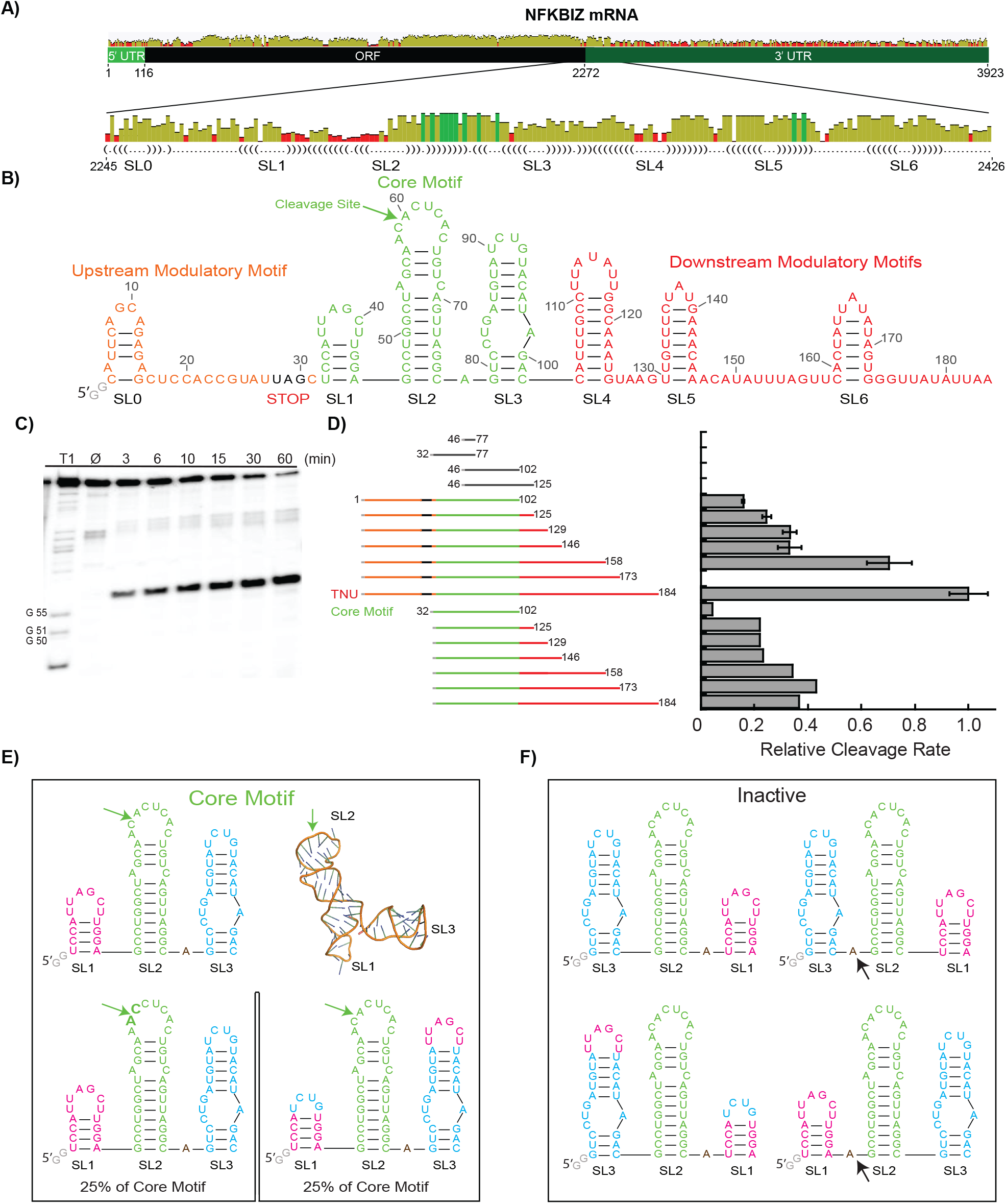
Fast and precise NFKBIZ cleavage results from the cooperation between a core cleavage domain and upstream and downstream modulatory domains. **A)** Conservation profile of NFKBIZ mRNA highlighting the 3′UTR region (NCBI RefSeq NM_031419.4). **B)** TNU construct with predicted secondary structure of its helical elements. The upstream motif (orange) and stop codon (black), the Regnase-1 cleavage core (green), and downstream modulatory motifs (red). The Regnase-1 cleavage site is indicated by the green arrow. Two guanosines were added to constructs to promote transcription (gray). **C)** 10% Denaturing gel showing *in vitro* cleavage with Regnase-1 (T1, RNase T1 ladder; Ø, no enzyme control). **D)** A set of inactive (black), and active TNU variants all containing the core cleavage motif (green). The cleavage efficiency of theses constructs relative to TNU is shown. **E)** Functional core motif constructs, and **F)** Inactive core motif variants, arrows note shift of adenosine from the left to the right side of SL2.

Further mapping of the TNU revealed that SL1, SL2, and SL3 were required to produce the precise cleavage pattern observed in **Figure 1C**. Only light and nonspecific cleavage occurring in the loop sequence region was observed with SL2 alone (fragment 46 to 77 of TNU) (**Figure S4.A**). Similarly, SL1/SL2 alone (fragment 32 to 77), SL2/SL3 alone (fragment 46 to 102) and SL2/SL3/SL4 (fragment 46 to 125) were inactive (**Figure 1D** and **S4.B-D**). The core motif (SL1/SL2/SL3 fragment 32 to 102) however was found to have significant and targeted cleavage that was nevertheless 25-fold slower than the TNU construct (**Figure 1D**). Adding only the upstream motif to the cleavage motif enhanced cleavage activity by 3.7-fold and adding the downstream stem-loops and single-stranded regions progressively increased activity to that of the full TNU construct (**Figure 1D**) indicating that both these stem-loops and the single-stranded regions help stabilize the pre-cleavage complex. The same series of downstream sequence extensions, but now lacking the upstream motif, showed considerably less cleavage activity (**Figure 1D**). Together, these experiments highlight the strong cooperativity between the upstream and downstream RNA elements, which are required for efficient RNA cleavage. It is noteworthy that the upstream domain exerts a strong modulatory effect, as it is easily imagined that active or stalled ribosomes could mask this region and thereby alter Regnase-1 activity. This would provide a mechanistic explanation for previous reports that the translational status^25^ of this mRNA strongly influences its regulation.

To probe the specificity of the core domain, we systematically rearranged its structural elements. Regnase-1 was found to recognize the stem regions rather than the loop sequences, despite the cleavage site being located within the SL2 loop. Swapping the two nucleotides flanking the cleavage site reduced activity only four-fold (**Figure 1E, lower left**), indicating that Regnase-1, unlike exonucleases such as RNase T1^28^, does not depend on a specific local RNA sequence. Similarly, exchanging the loop sequences of SL1 and SL3 slowed cleavage also by four-fold but did not abolish activity (**Figure 1E, lower right**). In contrast, swapping the entire SL1 and SL3 structures—or only their helical elements—eliminated cleavage (**Figure 1F, left**). These results suggest that the relative positioning of SL1 and SL3 helices is critical. Supporting this, repositioning the adenosine between SL2 and SL3 abolished activity in both wild-type and swapped contexts (**Figure 1F, right**). Collectively, these data indicate that Regnase-1 recognizes a core motif defined by the stems of SL1, SL2, and SL3, a structure we modeled using RNAComposer (**Figure 1E, top right**).

Previous studies have implicated the purine–purine–pyrimidine tri-loops found in SL5 and SL6 (**Figure 1B**) in Regnase-1–mediated repression of innate immune genes such as IL6 and NFKBIZ, and even suggested they may serve as direct cleavage targets^19,24,25^. However, replacing the UUUAUAUU loop of SL4 and the UAU tri-loops of SL5 and SL6 with a thermodynamically stable GAAA tetraloop^29,30^ altered Regnase-1 cleavage only modestly, indicating that these loops, while biologically important for NFKBIZ regulation^25^, are not strong determinants of Regnase-1-dependent cleavage. Consistent with this, Regnase-1 displayed only weak and nonspecific cleavage of SL5 and its destabilized variant (**Figure S5**). Taken together with the presence of zinc finger–like motifs in Regnase-1 outside its catalytic core^14^, these findings suggest that Regnase-1 stabilizes and positions its active site primarily through interactions with stem regions and adjacent single-stranded elements, rather than through strict loop-sequence recognition.

### Biologically cleavage is also precise and accelerated by subsequent uridylation

We next asked whether NFKBIZ is cleaved endogenously at the same position identified *in vitro* (**Figure 1**). Using MCF7 cells, we extracted total RNA, ligated adaptors to the 3′ ends, and performed a semi-nested RT–PCR targeting the predicted cleavage site. High-throughput sequencing revealed a predominant cleavage between C_59_ and A_60_, with the upstream C_59_ fragment heavily uridylated and accounting for 94.5% of total products (**Figure 2A**). The second most abundant product terminated at A_58_ (4.6% of reads) and was also uridylated. Additional products ending at A57 and C56, together with less frequent upstream and downstream cleavage fragments, represented only ∼0.9% of total reads but were likewise uridylated. These findings support the general conclusion that the primary cleavage between C_59_ and A_60_ by Regnase-1 is tightly coupled to uridylation, while the minor products likely reflect secondary processing events, such as partial degradation of the upstream fragment after Regnase-1 cleavage.

**Figure 2.**
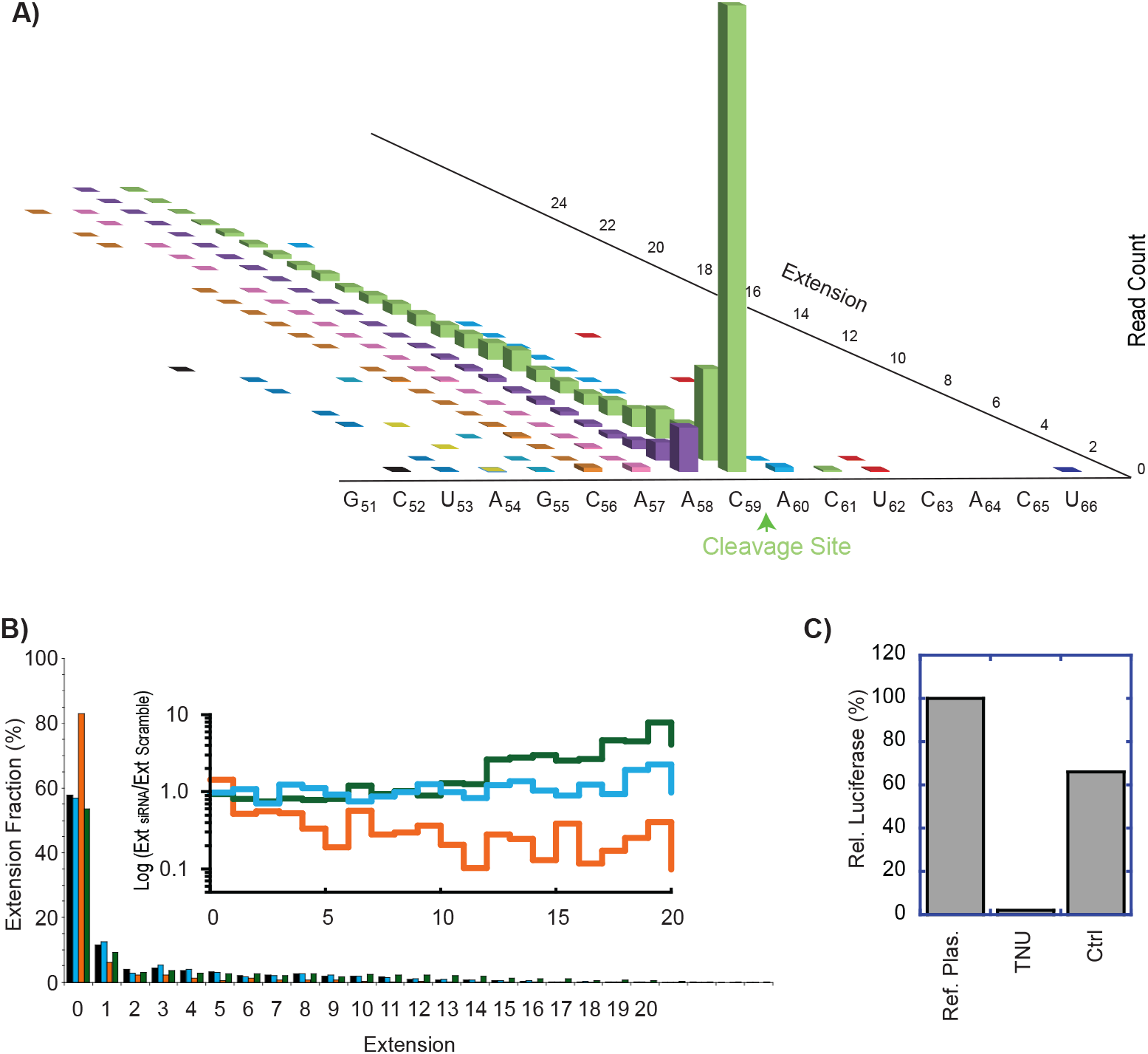
Regnase-1 cleavage site is uridylated by TUT7 and is subsequently degraded by DIS3L2. **A)** Sequencing of RT-PCR products identifies the NFKBIZ cleavage site and targeted uridylation. **B)** TUT7 siRNA knockdown (Orange data) decreases uridylation relative to a scramble control (Black data). TUT4 siRNA knockdowns remained comparatively flat (blue data). DIS3L2 siRNA knockdowns increased levels of longer uridylated products (Green data). Insert shows the log ratio of all three data sets relative to the siRNA scramble control (color codes as in the main histogram). **C)** Relative expression of luciferase for TNU.

Analysis of the uridylated products showed that the tails were heterogeneous, containing not only uridine but also adenosine (4.4%), guanosine (3.7%), and cytidine (0.5%) (**Figure S6**). This heterogeneous 3′ uridylation is characteristic of RNA targeted for degradation by the TUT– DIS3L2 pathway, previously implicated in mRNA, tRNA, rRNA, snRNA, snoRNA, and miRNA turnover^31–37^. We therefore tested whether TUT4/7 were responsible for uridylation. Consistent with this hypothesis, TUT7 and combined TUT4/7 knockdowns, but not TUT4 alone, reduced uridylation fractions by ∼5-fold for longer extensions relative to scrambled siRNA controls (**Figure 2B, orange data; Figure S7**). Correspondingly, untailed cleavage products at C_59_ accumulated in the TUT7 knockdown compared with a randomized siRNA control (**Figure 2B, inset**). Together, these results identify TUT7 activity as the primary source of the uridylation observed at the endogenous cleavage site

Because uridylation often directs transcripts to the highly processive 3′–5′ exonuclease DIS3L2^34,38,35^, we next tested whether DIS3L2 mediated turnover of uridylated NFKBIZ fragments. DIS3L2 knockdown increased the abundance of uridylated products longer than ∼12 nucleotides, with a two-to five-fold accumulation relative to scrambled controls (**Figure 2B, green data**). These findings being consistent with the established role of the TUT–DIS3L2 pathway as a general RNA surveillance mechanism degrading structured non-coding RNAs. To our knowledge, this represents the first direct evidence linking this pathway to targeted degradation of an mRNA central to immune regulation.

Importantly, *in vivo* Regnase-1 dependent cleavage and uridylation processing were not easily saturable: even high-level expression of artificial constructs resulted in precise cleavage and strong suppression of translation. In a luciferase reporter plasmid driven by a CMV promoter, fusing the TNU immediately downstream of the luciferase stop codon caused a ∼50-fold decrease in luciferase activity compared to the reference plasmid (**Figure 2C**). This strong repression was not seen in a control construct, which contained a downstream UTR fragment (nucleotides 749 to 930 of 3′ UTR), the same length as TNU (**Figure 2C**). Sequencing luciferase–TNU transcripts confirmed that cleavage again occurred precisely at the predicted site, and that the resulting fragments were again uridylated (**Figure S8**). These experiments demonstrate that the Regnase-1– TUT7–DIS3L2 pathway identified here does not require additional signals from the NFKBIZ ORF or distal 3’ UTR, but more importantly this regulatory process can effectively process and degrade transcripts when transcriptional levels are much higher than endogenous. This finding is consistent with our current understand of innate immunity where significant transcriptional upregulation can occur via the recruitment of transcription factors.

### B-cell driver mutations and viral RNA both stabilize the TNU segment against cleavage

A steady accumulation of literature points to the importance of NFKBIZ dysregulation upon the formation of certain types of B-Cell lymphomas and during viral infection. To explore the effect of B-cell lymphoma mutations we therefore measured *in vitro* cleavage rates for a panel of known B-cell lymphoma driver mutations^26^ in the TNU context (**Figure 3A)**. Such mutants in human patients typically only affect one NFKBIZ allele and its was striking that of the 15 mutants screened, 8 nearly significantly ablated Regnase-1 RNA cleavage and did so by altering the SL2 structure. A uridine insertion after A64, UG insertion after A64, the C65A, G67A, C69U and C65G mutations all would be predicted to destabilize the SL. Two deletion mutants targeted the right arm of SL2 and both significantly slowed cleavage (Δ67G-A70 and Δ65CU66). The location of these mutants was on the right arm of SL2, correlates well with the poor evolutionary conservation seen in the right arm of SL2 (**Figure 1A**). Notably a 11-nt deletion (Δ58A-G71) ablated targeted cleavage within SL2 completely but enhanced cleavage at G_100_ and C_102_, by 20-fold (**Figure S9**) indicating that the targeting to the core cleavage motif need not be absolute. In contrast, mutations in SL5 (U134C) and between SL5 and SL6 (U153A) showed no strong change in cleavage activity relative to TNU (**Figure 3A**).

**Figure 3.**
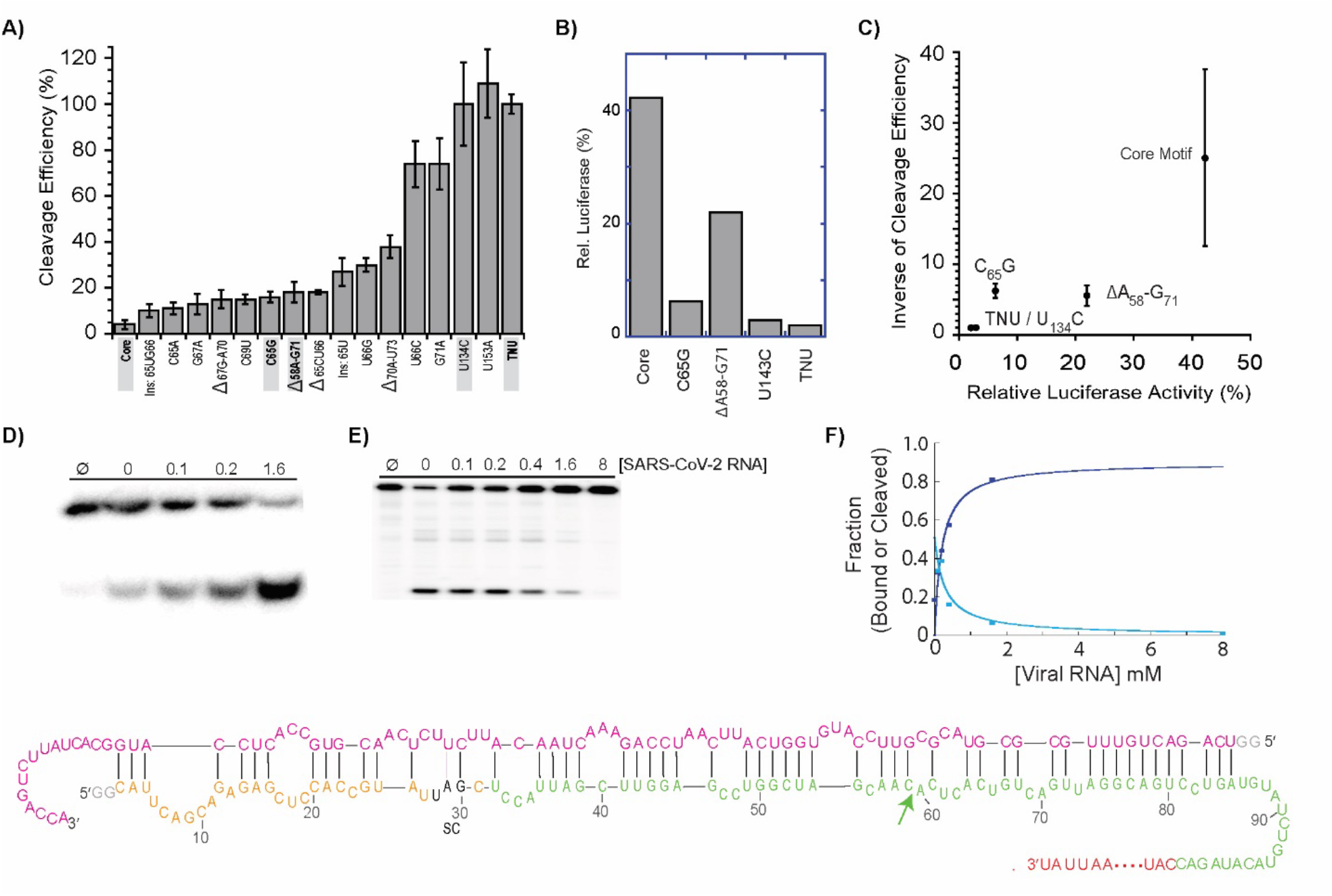
B-Cell lymphoma driver mutations and a segment of the SARS-CoV-2 genome both target the TNU region of NFKBIZ. **A)** Known B-cell NFKBIZ lymphoma mutants span a spectrum of *in vitro* cleavage rates. **B)** The luciferase expression of a subset of these mutants (names in gray boxes) when expressed in the TNU context. **C)** The inverse relationship between luciferase signal and *in vitro* mutant cleavage rates. **D)** TNU hybridizes to a SARS-CoV-2 fragment and **E)** can completely suppresses Regnase-1 cleavage in vitro. **F)** The fitted *K*_D_ = 201 ± 30 (dark blue curve) from panel D agrees well with the fitted inhibition constant of *K*_i_ = 282 ± 20 (light blue curve) from panel E. **G)** The SARS-CoV-2 genomic segment identified by Zhao et al. used to produce the gel shift with TNU.

Given this range of *in vitro* activities, we tested a representative set of these mutants *in vivo* using the TNU luciferase system. Interestingly, fusing only the core cleavage domain increased luciferase expression by 20-fold relative to the full TNU, consistent with the 25-fold reduction in cleavage efficiency measured *in vitro* (**Figure 3A and 3B**). Both C65G and the Δ58A-G71 deletion elevated luciferase expression, while the mutant U134C produced levels of luciferase similar to that of TNU (**Figure 3B**). Plotting the *in vivo* luciferase reporter data against the inverse of *in vitro* RNA cleavage rate (**Figure 3D**) further showed the inverse relationship between RNA cleavage rate and luciferase expression where higher mRNA cleavage results in less luciferase expression.

Remarkably, a recent study directly found that a segment of the SARs-CoV2 genome directly hybridizes to the TNU region of NFKBIZ^27^ and could cause a 3-fold increase in luciferase expression upon viral infection^27^. When we added the viral segment identified by Zaho et al and titrated it with respect to a ^32^P radiolabeled TNU substrate we could observe both a complete native gel shift (**Figure 3D**) and corresponding suppression of Regnase-1 dependent cleavage (**Figure 3E**), with the *K*_D_ and *K*_I_ agreeing well (**Figure 3F**). Strikingly, the rescue they achieved *in vivo* by mutating the viral sequence element (**Figure 3G**) in our hands reduced RNA cleavage by two-fold. Thus, in addition to the B-cell lymphoma data just discussed the TNU / Regnase-1 cleavage-dependent regulatory mechanism appears to be actively suppressed during viral infection by the formation of bimolecular secondary structure which prevents recognition by Regnase-1.

## DISCUSSION

We have defined a rapid and targeted cleavage mechanism that maintains NFKBIZ expression at very low levels under normal cellular conditions. Regnase-1 cleavage of the NFKBIZ 3′UTR is tightly coupled to TUT7-dependent uridylation and subsequent DIS3L2-mediated degradation, providing for the first-time mechanistic insight into the rapid turnover of the NFKBIZ mRNA as summarized in **Figure 4**. In both lymphoma and viral infection, this pathway can be bypassed: driver mutations alter the TNU structure, while viral RNAs such as SARS-CoV-2 mask it, in each case elevating NFKBIZ levels presumably dysregulate the normal NFKBIZ response to such insults.

**Figure 4.**
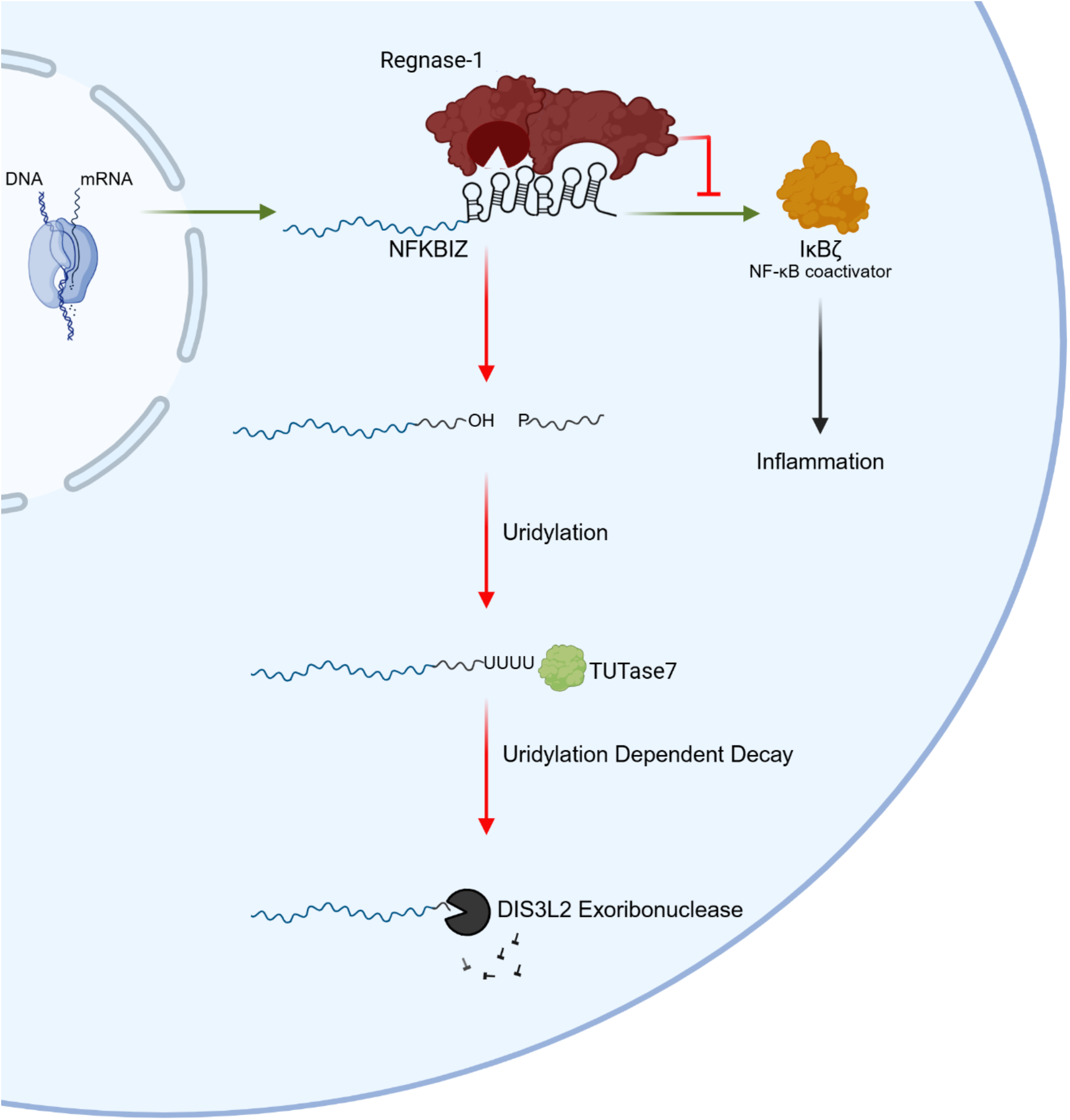
The Regnase-1 dependent cleavage model for NFKBIZ regulation. RNAs recognized by Regnase-1 such as the TNU element in NFKBIZ can be rapidly cleaved triggering uridylation by TUT7 and subsequent degradation of the ORF by DIS3L2. Suppression of RNA cleavage by RNA binding/remodeling factors, mutation in certain cancers and hybridization of viral genomic RNA to the cleavage site all can release this regulatory inhibition.

Previous models of Regnase-1 activity invoked rapid transcript decay but lacked an explicit molecular mechanism. Mino et al. proposed that Regnase-1 targets pyrimidine–purine–pyrimidine stem-loops within SL5 and SL6 of the TNU via UPF1-mediated remodeling, and that cleavage is suppressed during chronic infection by Roquin binding to these elements^25^. Our findings reinterpret this model: Roquin binding to SL5 and SL6 likely prevents these and adjacent elements from engaging Regnase-1, thereby indirectly blocking cleavage and thus allowing a transition into chronic infection. Consistent with this model B-cell lymphoma mutations are known where Regnase-1 itself is mutated.

We further demonstrated that the upstream motif strongly enhances Regnase-1 cleavage efficiency, providing a means to integrate translational status into this regulatory circuit. Ribosomes stalled in the 3′ ORF or at the stop codon would be expected to obstruct the upstream motif, further attenuating Regnase-1 access. In combination with Roquin and remodeling factors such as UPF1, this arrangement will allow rapid, and highly context-dependent modulation of NFKBIZ levels via the Regnase-1–TUT7–DIS3L2 pathway characterized here.

As Regnase-1 regulates many innate immune genes containing similar stem-loop structures to those found in the NFKBIZ UTR, our results suggest that additional cancers and viruses may exploit mutation and RNA binding strategies, respectively, to help evade the innate immune response. Thus, the NFKBIZ TNU–Regnase-1 type cleavage mechanism characterized here for NFKBIZ may occur for other innate immune response genes and may represent a general vulnerability in immune regulation and thus be a potential target for therapeutic intervention.

## MATERIALS AND METHODS

### Protein expression and purification

The cDNA encoding Regnase-1 (NP_001310479.1; 1–599 aa) was fused to a 6xHis tag in the pET-30 expression vector (Sigma-Aldrich). pET-Reg1-HIS was transformed into Rosetta (DE3) Competent Cells (Novagen) and grown until they reached and optical density of OD_600_ of 0.6. Regnase-1 gene expression was induced by adding 0.4 mM Isopropyl b-D-1-thiogalactopyranoside (Sigma-Aldrich) for 4 hours, and cells were harvested/lysed by 3 rounds of flash freezing (liquid nitrogen) and French press. Crude cell extract was pelleted, and the supernatant was batch bound to Ni-NTA beads pre-equilibrated with binding buffer (PBS, pH 7.4, 10 mM imidazole, 1 mM PMSF) for 1 hour at 4°C, followed by transfer to a column. Washing and elution were performed with the same binding buffer supplemented with 50 mM and 250 mM imidazole, respectively. Eluted protein was dialyzed into storage buffer (50 mM Tris-Base, pH 7.4, 150 mM NaCl, 1 mM EDTA, 1 mM DTT, 50% Glycerol) overnight at 4°C. Protein was then aliquoted and flash frozen with liquid nitrogen and kept in –80°C until use.

### *In vitro* transcription of wild-type TNU, and NFKBIZ 3′ UTR RNA variants

A fragment of the human NFKBIZ 3′ UTR (NCBI RefSeq: NM_031419.4; nucleotides 2244– 2426, TNU) and its mutant variants were synthesized as single-stranded DNA oligonucleotides (Integrated DNA Technologies, Coralville, IA). Double-stranded DNA templates were generated by primer extension and transcribed in vitro using T7 RNA polymerase. To enhance transcription efficiency, a T7 promoter followed by two additional guanosines was incorporated at the 5′ end of each DNA template; as a result, all RNA variants contained two extra G residues at their 5′ termini. Transcription reactions were performed using 0.5 µM dsDNA template, 8 mM GTP, 2 mM UTP, 5 mM each of CTP and ATP, and T7 transcription buffer (40 mM Tris-HCl, pH 8.0 at 37°C, 26 mM MgCl_2_, 2.5 mM spermidine, 10 mM DTT, and 0.01% Triton X-100), supplemented with T7 RNA polymerase. Reactions were incubated at 37°C for 1.5 h. RNA products were denatured in 50% formamide and 10 mM EDTA at 95°C for 5 min, separated by 5% denaturing PAGE gel, and visualized by UV shadowing. Gel slices containing the target RNA were excised and incubated overnight in 300 mM NaCl at 4°C with rotation. Eluted RNA was precipitated with 2.5 equivalents of ethanol, resuspended in RNase-free water, and quantified by absorbance at 260 nm using a NanoDrop 2000 spectrophotometer (Thermo Fisher Scientific) and IDT ‘ s online extinction coefficient calculator.

### RNA 5′ end dephosphorylation and radiolabeling

Purified RNA (5 µM) was treated with calf intestinal alkaline phosphatase (CIP; New England BioLabs) in a reaction buffer containing 50 mM potassium acetate, 20 mM Tris-acetate, 10 mM magnesium acetate, and 100 µg/ml recombinant albumin (pH 7.9 at 25°C), at a final concentration of 1 U/μl. Dephosphorylation was performed at 37°C for 30 min, followed by phenol-chloroform extraction and ethanol precipitation. CIP-treated RNA was then 5′-end labeled using T4 polynucleotide kinase (PNK; New England BioLabs) in the presence of 0.1 µM [γ-32P] ATP, 10 mM MgCl_2_, 5 mM DTT, and 70 mM Tris-HCl (pH 7.6 at 25°C), with 1 U/µl PNK. Labeling reactions were incubated at 37°C for 30 min, then heat-inactivated and purified by phenol-chloroform extraction and ethanol precipitation.

### *In vitro* Regnase-1 cleavage assays

5′-radiolabeled RNA substrates were incubated with recombinant Regnase-1 at 37°C for varying time points in cleavage buffer containing 25 mM HEPES, 50 mM potassium acetate, 5 mM DTT, and 5 mM magnesium acetate, 1 U of SUPERase (Thermo Fisher), pH 7.5. Reactions were terminated by the addition of 50% formamide and 10 mM EDTA, followed by heat denaturation at 95°C for 5 min. Cleavage products were resolved on 10% denaturing PAGE and visualized using a GE Healthcare Amersham Typhoon phosphor imager.

To generate an alkaline hydrolysis ladder, 100 pM of 5′-radiolabeled RNA was incubated in 50 mM sodium bicarbonate (NaHCO_3_) at 90°C for 5 min. Reactions were stopped with 100 mM Tris-HCl and an equal volume of loading buffer containing 90% formamide and 20 mM EDTA, followed by denaturation at 95°C for 5 min. Cleavage products were resolved on 10% denaturing polyacrylamide sequencing gels and visualized using a GE Healthcare Amersham Typhoon phosphor imager.

100 pM of 5′-radiolabeled RNA was incubated in a reaction mixture containing 6 M urea, 20 mM sodium citrate (pH 5.0 at 25°C), and 0.2 U RNase T1 (Thermo Fisher Scientific). Reactions were carried out at 50°C for 10 min following the addition of loading buffer containing 90% formamide and 20 mM EDTA. Samples were denatured at 95°C for 5 min, resolved on 10% denaturing polyacrylamide sequencing gels, and visualized using a GE Healthcare Amersham Typhoon phosphor imager.

### siRNA/plasmid transfections, Luciferase assays and total RNA extractions

30pmol of siRNA targeting TUT4 (ThermoFisher, siRNA ID: 259168), TUT7 (ThermoFisher, siRNA ID: siRNA ID:), DIS3L2 (ThermoFisher, siRNA ID: 126074), or control (ThermoFisher, SilencerTM Select Negative Control No.1 siRNA) was transfected into MCF7 cells using LipofectamineTM 3000 reagent (Invitrogen). 48-hours following transfection, samples were harvested with TriZol. Variants of the NFKBIZ 3′ UTR were synthesized as gBlocks (Integrated DNA Technologies) and cloned into the pGL3-Promoter (Promega) downstream of the luciferase stop codon. 0.5 μg of these constructs were co-transfected with 0.1 μg of pRL-TK (Promega) into MCF-7 cells using LipofectamineTM 3000 (Invitrogen), and cells were incubated for 24-hours transfection media. Lysis and dual luciferase detection were performed according to manufacturer’s instructions (Promega). Firefly luciferase signals were normalized to the Renilla luciferase transfection control, and transfections were done in biological triplicates. Total RNA was extracted using TRIzol reagent on cells that were 80–90% confluence, according to the manufacture’s protocol (ThermoFisher). Prior to subsequent assays RNA integrity was assessed by agarose gel electrophoresis and quantified via NanoDrop.

### Adenylated Oligo Ligation and cDNA Amplification

1.5 mg of Total RNA from MCF7 cells was ligated to an adenylated DNA Oligo (IDT) with the sequence /5rApp/GAAGAGCCTACGACGA/3ddC/ (adapter at 20 μM, 50 mM HEPE pH 8.3, 10 mM MgCl2, 3.3 mM DTT, 10% PEG 4000, 10% DMSO) using (1 U) T4 RNA ligase (NEB) for 90 min at room temperature. Excess adenylated oligo was removed by passing the ligation through a 100 kDa spin column twice. Adenylated RNAs were reverse transcribed using a 20 µM of primer complementary to the adenylated oligo using Maxima Reverse Transcriptase according to the manufacturer’s instructions. Again, excess reverse transcription primer was washed of by spinning down reaction down 100 kDa spin column twice. An outer PCR reaction having forward primer was performed for 20 cycles, and was then diluted 100-fold into an inner PCR reaction containing forward primer for a further 35 cycles. Bands from a 2 % agarose gel were excised and purified (QIAquick Gel Extraction Kit) and submitted for Amplizon Azenta HTS sequencing (PCR reaction conditions).

## REFERENCES

1. Feng Y, Chen Z, Xu Y, Han Y, Jia X, Wang Z, et al. The central inflammatory regulator IκBζ: induction, regulation and physiological functions. Front Immunol. 2023 June 12;14:1188253.

2. Kombe Kombe AJ, Fotoohabadi L, Gerasimova Y, Nanduri R, Lama Tamang P, Kandala M, et al. The Role of Inflammation in the Pathogenesis of Viral Respiratory Infections. Microorganisms. 2024 Dec 7;12(12):2526.

3. Wu B, Sodji QH, Oyelere AK. Inflammation, Fibrosis and Cancer: Mechanisms, Therapeutic Options and Challenges. Cancers. 2022 Jan 22;14(3):552.

4. Hirano T. IL-6 in inflammation, autoimmunity and cancer. Int Immunol. 2021 Mar 1;33(3):127–48.

5. L. Kiss A. Inflammation in Focus: The Beginning and the End. Pathol Oncol Res. 2022 Jan 4;27:1610136.

6. Takeuchi O, Akira S. Pattern Recognition Receptors and Inflammation. Cell. 2010 Mar;140(6):805–20.

7. Khabar KSA. Post-transcriptional control during chronic inflammation and cancer: a focus on AU-rich elements. Cell Mol Life Sci. 2010 Sept;67(17):2937–55.

8. Uehata T, Akira S. mRNA degradation by the endoribonuclease Regnase-1/ZC3H12a/MCPIP-1. Biochim Biophys Acta BBA - Gene Regul Mech. 2013 June;1829(6–7):708–13.

9. Hodson DJ, Janas ML, Galloway A, Bell SE, Andrews S, Li CM, et al. Deletion of the RNA-binding proteins ZFP36L1 and ZFP36L2 leads to perturbed thymic development and T lymphoblastic leukemia. Nat Immunol. 2010 Aug;11(8):717–24.

10. Wang X, Tang G, Liu Y, Zhang L, Chen B, Han Y, et al. The role of IL-6 in coronavirus, especially in COVID-19. Front Pharmacol. 2022 Nov 23;13:1033674.

11. Law HKW, Cheung CY, Ng HY, Sia SF, Chan YO, Luk W, et al. Chemokine up-regulation in SARS-coronavirus–infected, monocyte-derived human dendritic cells. Blood. 2005 Oct 1;106(7):2366–74.

12. Li W, Li F, Zhang X, Lin HK, Xu C. Insights into the post-translational modification and its emerging role in shaping the tumor microenvironment. Signal Transduct Target Ther. 2021 Dec 20;6(1):422.

13. Pararajalingam P, Coyle KM, Arthur SE, Thomas N, Alcaide M, Meissner B, et al. Coding and noncoding drivers of mantle cell lymphoma identified through exome and genome sequencing.

14. Liang J, Wang J, Azfer A, Song W, Tromp G, Kolattukudy PE, et al. A Novel CCCH-Zinc Finger Protein Family Regulates Proinflammatory Activation of Macrophages. J Biol Chem. 2008 Mar;283(10):6337–46.

15. Matsushita K, Takeuchi O, Standley DM, Kumagai Y, Kawagoe T, Miyake T, et al. Zc3h12a is an RNase essential for controlling immune responses by regulating mRNA decay. Nature. 2009 Apr;458(7242):1185–90.

16. Uehata T, Iwasaki H, Vandenbon A, Matsushita K, Hernandez-Cuellar E, Kuniyoshi K, et al. Malt1-Induced Cleavage of Regnase-1 in CD4+ Helper T Cells Regulates Immune Activation. Cell. 2013 May;153(5):1036–49.

17. Iwasaki H, Takeuchi O, Teraguchi S, Matsushita K, Uehata T, Kuniyoshi K, et al. The IκB kinase complex regulates the stability of cytokine-encoding mRNA induced by TLR–IL-1R by controlling degradation of regnase-1. Nat Immunol. 2011 Dec;12(12):1167–75.

18. Musson R, Szukała W, Jura J. MCPIP1 RNase and Its Multifaceted Role. Int J Mol Sci. 2020 Sept 29;21(19):7183.

19. Behrens G, Winzen R, Rehage N, Dörrie A, Barsch M, Hoffmann A, et al. A translational silencing function of MCPIP1/Regnase-1 specified by the target site context. Nucleic Acids Res. 2018 May 4;46(8):4256–70.

20. Li H, Wang TT. MCPIP1/regnase-I inhibits simian immunodeficiency virus and is not counteracted by Vpx. J Gen Virol. 2016 July 1;97(7):1693–8.

21. Lin RJ, Chu JS, Chien HL, Tseng CH, Ko PC, Mei YY, et al. MCPIP1 Suppresses Hepatitis C Virus Replication and Negatively Regulates Virus-Induced Proinflammatory Cytokine Responses. J Immunol. 2014 Oct 15;193(8):4159–68.

22. Mizgalska D, Węgrzyn P, Murzyn K, Kasza A, Koj A, Jura J, et al. Interleukin-1-inducible MCPIP protein has structural and functional properties of RNase and participates in degradation of IL-1β mRNA. FEBS J. 2009 Dec;276(24):7386–99.

23. Yamazaki S. The Nuclear NF-κB Regulator IκBζ: Updates on Its Molecular Functions and Pathophysiological Roles. Cells. 2024 Aug 31;13(17):1467.

24. Mino T, Iwai N, Endo M, Inoue K, Akaki K, Hia F, et al. Translation-dependent unwinding of stem–loops by UPF1 licenses Regnase-1 to degrade inflammatory mRNAs. Nucleic Acids Res. 2019 July 22;gkz628.

25. Mino T, Murakawa Y, Fukao A, Vandenbon A, Wessels HH, Ori D, et al. Regnase-1 and Roquin Regulate a Common Element in Inflammatory mRNAs by Spatiotemporally Distinct Mechanisms. Cell. 2015 May;161(5):1058–73.

26. Arthur SE, Jiang A, Grande BM, Alcaide M, Cojocaru R, Rushton CK, et al. Genome-wide discovery of somatic regulatory variants in diffuse large B-cell lymphoma. Nat Commun. 2018 Oct 1;9(1):4001.

27. Zhao H, Cai Z, Rao J, Wu D, Ji L, Ye R, et al. SARS-CoV-2 RNA stabilizes host mRNAs to elicit immunopathogenesis. Mol Cell. 2024 Feb;84(3):490-505.e9.

28. Pace CN, Heinemann U, Hahn U, Saenger W. Ribonuclease T1: Structure, Function, and Stability. Angew Chem Int Ed Engl. 1991 Apr;30(4):343–60.

29. Jucker FM, Heus HA, Yip PF, Moors EHM, Pardi A. A Network of Heterogeneous Hydrogen Bonds in GNRA Tetraloops. J Mol Biol. 1996 Dec;264(5):968–80.

30. Menger M, Eckstein F, Porschke D. Dynamics of the RNA Hairpin GNRA Tetraloop. Biochemistry. 2000 Apr 1;39(15):4500–7.

31. Lim J, Ha M, Chang H, Kwon SC, Simanshu DK, Patel DJ, et al. Uridylation by TUT4 and TUT7 Marks mRNA for Degradation. Cell. 2014 Dec;159(6):1365–76.

32. Lim J, Ha M, Chang H, Kwon SC, Simanshu DK, Patel DJ, et al. Uridylation by TUT4 and TUT7 Marks mRNA for Degradation. Cell. 2014 Dec;159(6):1365–76.

33. Thornton JE, D. P, Jing L, Sjekloca L, Lin S, Grossi E, et al. Selective microRNA uridylation by Zcchc6 (TUT7) and Zcchc11 (TUT4). Nucleic Acids Res. 2014 Oct 13;42(18):11777–91.

34. Ustianenko D, Pasulka J, Feketova Z, Bednarik L, Zigackova D, Fortova A, et al. TUT-DIS3L2 is a mammalian surveillance pathway for aberrant structured non-coding RNAs. EMBO J. 2016 Oct 17;35(20):2179–91.

35. Yang A, Shao TJ, Bofill-De Ros X, Lian C, Villanueva P, Dai L, et al. AGO-bound mature miRNAs are oligouridylated by TUTs and subsequently degraded by DIS3L2. Nat Commun. 2020 June 2;11(1):2765.

36. Medhi R, Price J, Furlan G, Gorges B, Sapetschnig A, Miska EA. RNA uridyl transferases TUT4/7 differentially regulate miRNA variants depending on the cancer cell type.

37. Zhang P, Frederick MI, Heinemann IU. Terminal Uridylyltransferases TUT4/7 Regulate microRNA and mRNA Homeostasis. Cells. 2022 Nov 23;11(23):3742.

38. Da Costa PJ, Menezes J, Saramago M, García-Moreno JF, Santos HA, Gama-Carvalho M, et al. A role for DIS3L2 over natural nonsense-mediated mRNA decay targets in human cells. Biochem Biophys Res Commun. 2019 Oct;518(4):664–71.

