## Supplemental Figures for "Regulation of NFKBIZ by precise Regnase-1 endoribonuclease cleavage and subsequent uridylation"

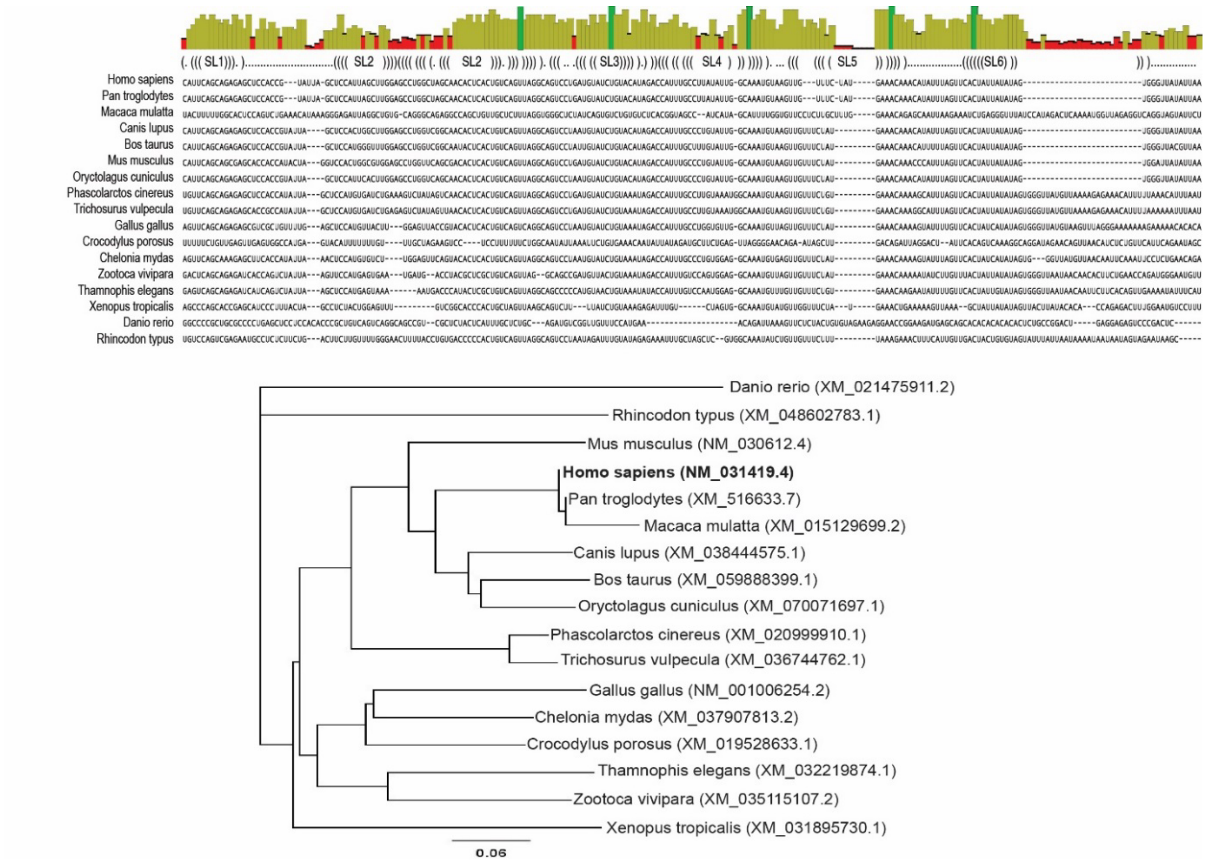


**Figure S1**. Conservation pattern of NFKBIZ alignment from human (*Homo sapiens*) to zebrafish (*Danio rerio*).

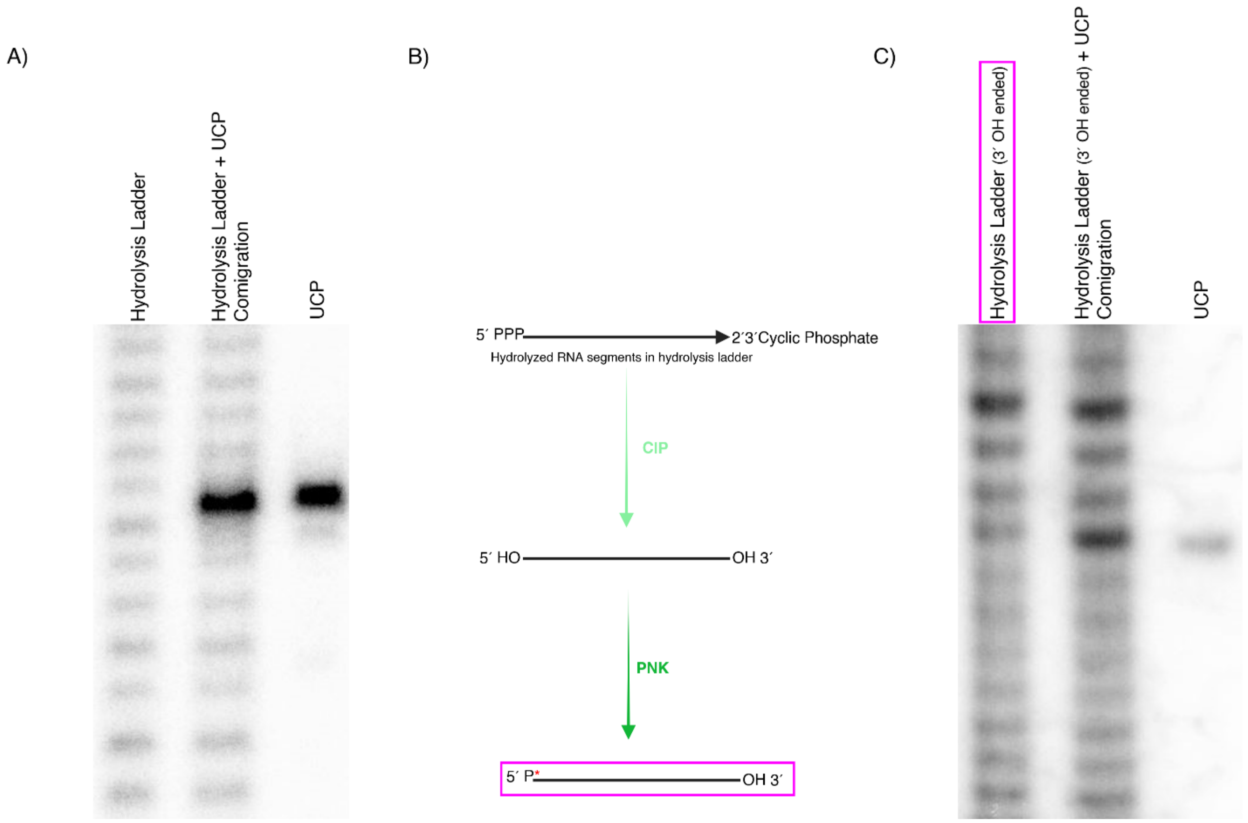


**Figure S2. The Upstream Cleavage Product (UCP) generated by Regnase-1 carries a 3′ hydroxyl terminus, as determined by comigration analysis.**
**A)** UCP did not comigrate with an alkaline-hydrolyzed RNA ladder, which contains fragments terminating with 2′,3′-cyclic phosphates. The hydrolysis ladder (lane 1) and the UCP (lane 3) displayed distinct migration patterns, and their mixture (lane 2) confirmed the difference. **B)** Conversion of 2′,3′-cyclic phosphate termini to 3′ hydroxyl termini: Alkaline-hydrolyzed RNA was treated with calf intestinal phosphatase (CIP) followed by polynucleotide kinase (PNK) to generate a modified ladder bearing 3′ hydroxyl groups and labeled at their 5' ends as before. **C)** This new modified ladder (lane 1) comigrated with the UCP (lane 3), as shown by their mixture (lane 2).

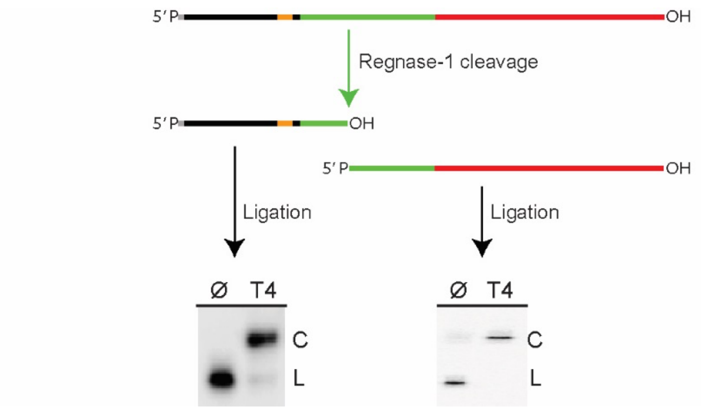


**Figure S3. T4 RNA ligation also demonstrates a 3’ hydroxyl on the upstream fragment and a 5’ phosphate on the downstream fragment.** Schematic of Regnase-1 cleavage products and subsequent ligation assay. Upstream (left) and downstream (right) cleavage fragments were gel purified and subjected to T4 RNA ligase treatment, generating circular (C) from the initial linear (L) products.

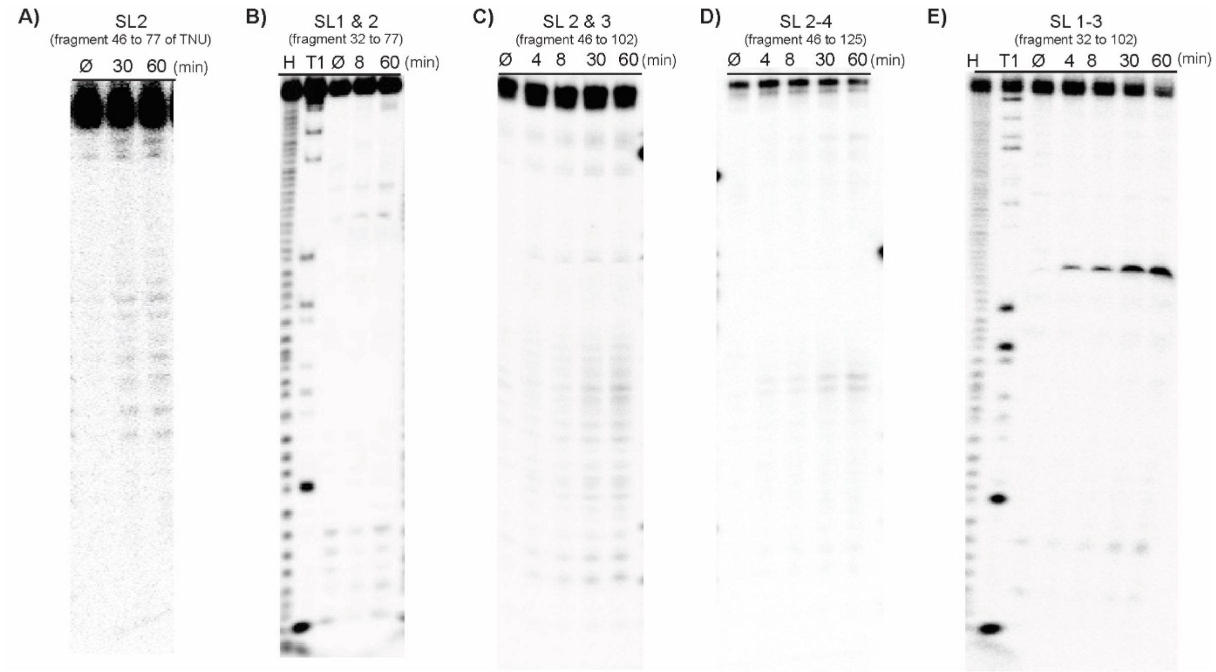


**Figure S4: Target degradation occurs when SL1,2 &3 are present but not its subcomponents.** Inactive truncated TNU motifs and the core motif with a defined cleavage site resolved on a 10% denaturing PAGE gel (1.5 h, 58 W).

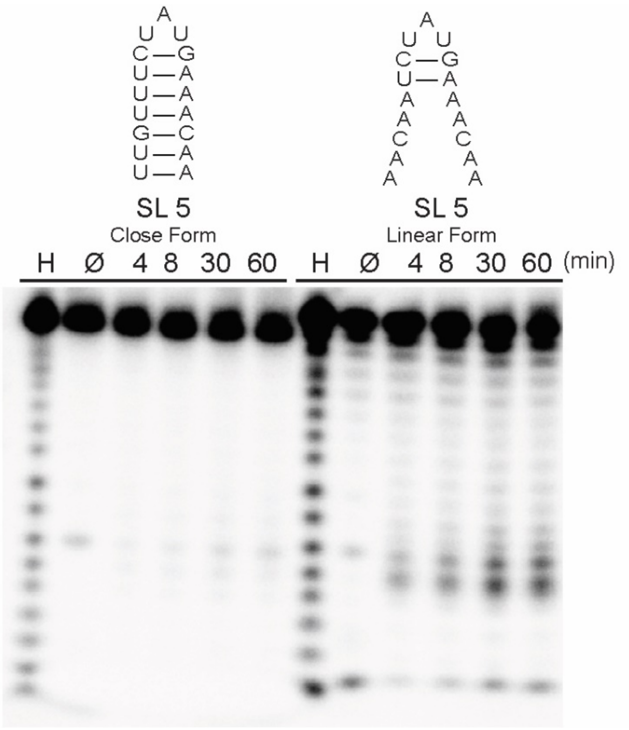


**Figure S5. Regnase-1 cleavage of closed versus open SL5.** Wild-type SL5 shows very slow degradation whereas the linear form undergoes extensive non-specific degradation by Regnase-1. Constructs adapted from: Mino, T. et al. Translation-dependent unwinding of stem-loops by UPF1 licenses Regnase-1 to degrade inflammatory mRNAs. Nucleic Acid Research, (2019).

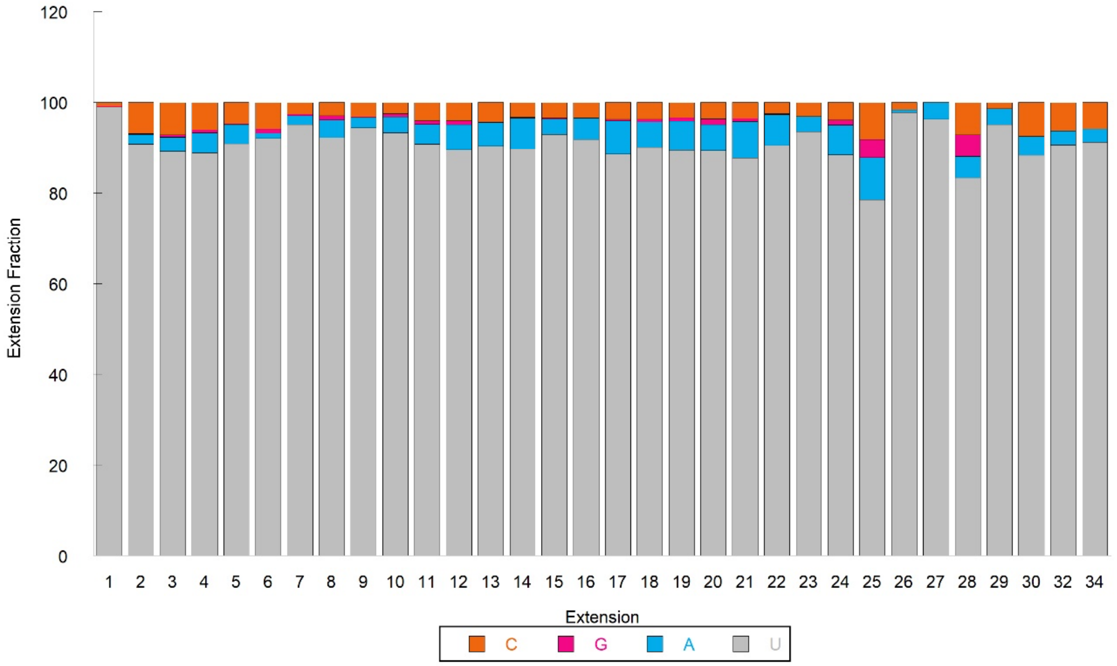


**Figure S6: Uridinylation products are not homogenous U sequences after the first addition.** Nucleotide composition of extension products of the endogenous sample at the dominant cleavage site at $C_{59}$.

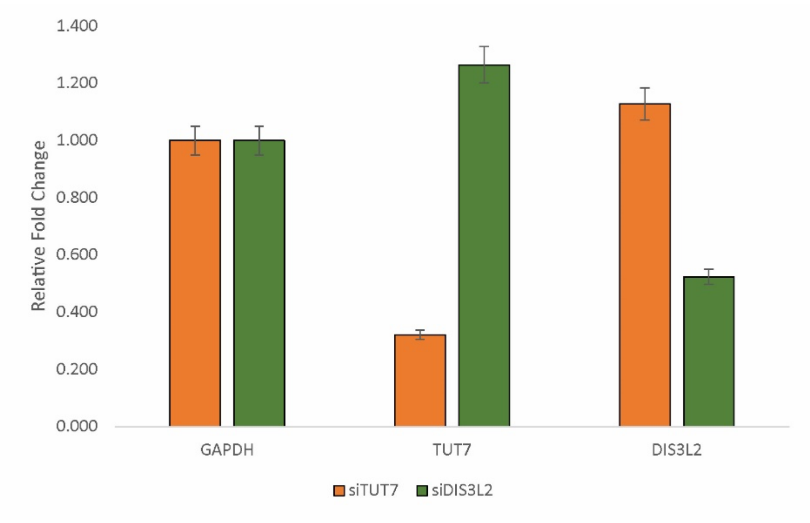


**Figure S7.** qRT-PCR results for siRNA libraries, normalized to GAPDH of the scrambled library.

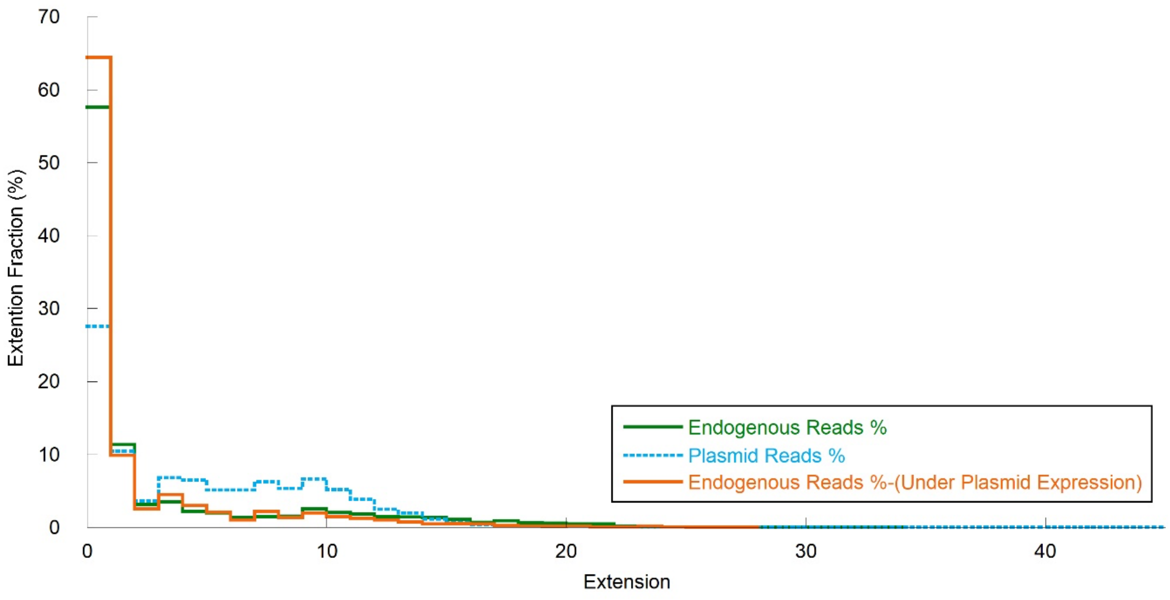


**Figure S8:** Tail distribution at the dominant cleavage site for endogenous NFKBIZ 3′UTR, TNU inserted downstream of a luciferase plasmid, endogenous NFKBIZ 3'UTR under plasmid expression.

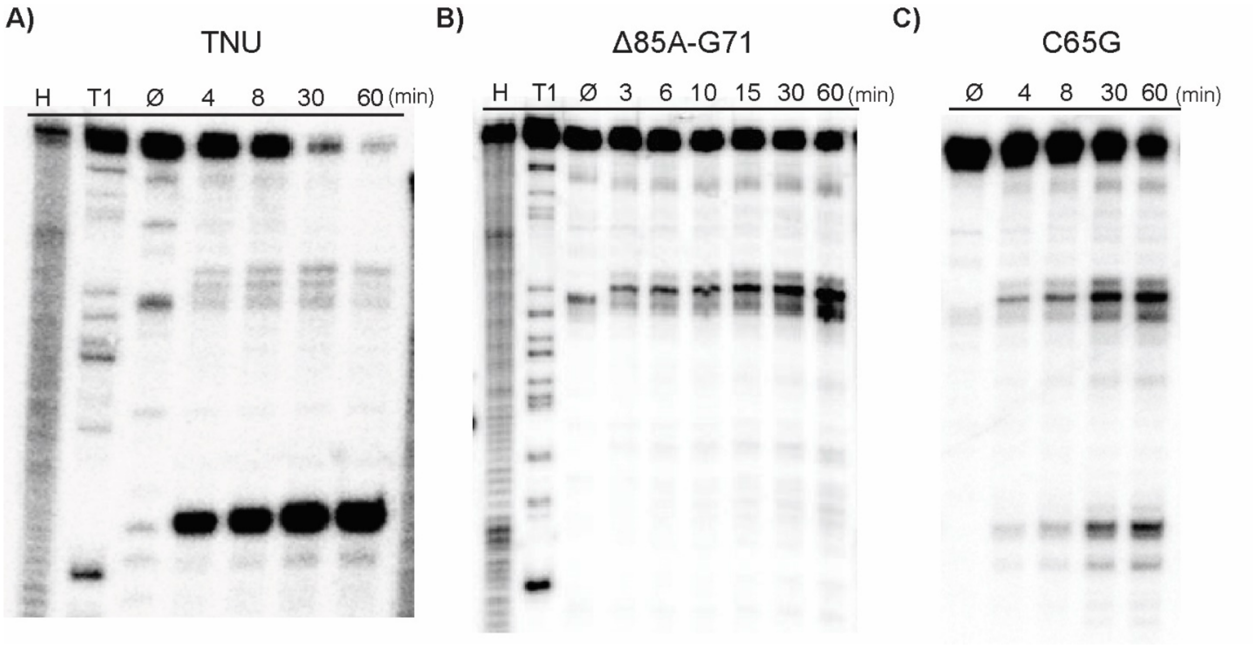


**Figure S9. Cleavage patterns of Regnase-1 for two cancer-associated mutants.** Regnase-1 cleavage of TNU. **B)** An 11-nt deletion removes the dominant cleavage site, redirecting cleavage to a minor site within TNU. **C)** The C65G mutation substantially reduces cleavage and partially shifts it from the dominant to secondary sites.
